# Multicenter Study of the Burden of Multidrug-Resistant Bacteria in the Etiology of Infected Diabetic Foot Ulcers

**DOI:** 10.1101/625012

**Authors:** Adeyemi Temitayo Adeyemo, Babatope A. Kolawole, Vincent Olubunmi Rotimi, Aaron Oladipo Aboderin

## Abstract

**Background:** Infected diabetic foot ulcer (IDFU) is a public health issue and a leading cause of non-traumatic limb amputation. Very few published data on IDFU is available in most West African countries. The objective of this study was to investigate the etiological agents of IDFU and the challenge of antibacterial drug resistance in the management of infections.

**Methods:** This was a prospective cross-sectional hospital-based study involving three tertiary healthcare facilities. Consecutive eligible patients presenting in the facilities were recruited. Tissue biopsies and/or aspirates were collected and cultured on a set of selective and non-selective media and incubated in appropriate atmospheric conditions for 24 to 72 hours. Isolates were identified by established standard methods. Antibiotic susceptibility testing was performed using modified Kirby-Bauer disc diffusion method. Specific resistance determinants were investigated by polymerase chain reaction-based protocols. Data analysis was done with SPSS version 20.

**Results:** Ninety patients with clinical diagnosis of DFI were studied between July 2016 and April 2017. A total of 218 microorganisms were isolated, comprising 129 (59.2%) Gram-negative bacilli (GNB), 59 (27.1%) Gram-positive cocci (GPC) and 29 (13.2%) anaerobic bacteria. The top five facultative/aerobic bacteria encountered were: *Staphylococcus aureus*, *Escherichia coli*, *Pseudomonas aeruginosa*, *Klebsiella pneumoniae* and *Citrobacter* spp. representing 41 (18.8%), 23 (10.5%), 20 (9.2%), 19 (8.7%) and 19 (8.7%) isolates, in that order, respectively. The commonest anaerobes were *Bacteroides* spp., and *Peptostreptococcus anaerobius* which accounted for 7 (24.1%) and 6 (20.7%), respectively. Of the 93 IDFU cases, 74 (80%) were infected by multidrug-resistant (MDR) bacteria predominantly methicillin-resistant *S. aureus*, extended-spectrum β-lactamase-producing GNB, mainly of the CTX-M variety. Only 4 (3.1%) GNB were carbapenemase-producers encoded by *bla*_VIM_. Factors associated with presence of MDR bacteria were peripheral neuropathy (r= 4.05, *P*= 0.042) and duration of foot infection >1 month(r= 7.63, *P*= 0.015).

**Conclusions:** MDR facultative/aerobic bacteria are overrepresented amongst agents causing IDFU. A relatively low proportion of the etiological agents were anaerobic bacteria. This finding should help formulate empirical therapeutic options for managing IDFU. Furthermore, drastic reduction in inappropriate use of cocktail of antibiotics for IDFUs is advocated to combat infection by MDR bacteria in these patients.

## Introduction

Infected diabetic foot ulcer (IDFU) is associated with inflammation or purulence occurring in a site below the ankle in a person with diabetes mellitus (DM).^1^ It is a major global public health with substantial medical, socio-economic and psychological burden. IDFU is one of the most common diabetes-related infections in clinical practice, and a common indication for hospital admission.^2^ At 7.2% (95%CI: 5.1-9.3%) and higher than the global prevalence of 6.3% (95%CI: 5.4-7.3%), Africa has the second highest global prevalence of diabetic foot ulcer, a precursor of IDFU.^3^ Paradoxically, foot infections are most common and lethal in Africa than elsewhere globally.^4^ Between 25-60% of diabetic patients with background foot ulcer will develop IDFU which remains a major reason for non-traumatic amputation of the lower limbs.^5^

Wide varieties of organisms, including anaerobic bacteria, have been implicated in the etiology of IDFU depending on severity of infection and time from onset to presentation at the healthcare facility. Advanced IDFUs with features of sepsis at admission usually harbor anaerobic pathogens.^2^ Emergence and current global threat of antimicrobial resistance in the face of dwindling antibiotics in the pipeline has added a new twist to the burden of IDFU.^6^ Increasing involvement of multi-drug resistant organisms in diabetic patients with infected foot ulcers has significantly reduced antibiotic treatment options, thus posing a serious challenge in resource-constrained low- and middle-income countries where access to antimicrobial drugs is of grave concern.^7^ A recent systematic review and meta-analysis on the global burden of diabetic foot ulceration in Cameroon, a West African country, has concluded that paucity of data is a bane on the strategy for treatment and prevention of foot infections in diabetic patients.^3^ Thus, our study was designed to determine the prevalent bacteria involved in IDFUs, assess the burden of MDR bacteria among the isolates and evaluate the associated risk factors.

## Materials and Methods

### Patients

This prospective cross-sectional hospital-based multicenter study was carried out at the Obafemi Awolowo University Teaching Hospitals Complex, Ile-Ife (Ife Hospital Unit, Wesley Guild Hospital, Ilesa and Ladoke Akintola University of Technology Teaching Hospital, Osogbo between July 2016 and April 2017. All consecutive diabetic patients with foot infections meeting the criteria for the diagnosis of IDFU, seen and managed at these hospitals, were recruited into the study. They were clinically assessed and foot lesions graded according to diabetic foot infection severity classification system issued by the Infectious Disease Society of America.^2^ Only non-duplicate patients and samples were studied. Ethical approvals for this study were granted by the Ethics and Research Committees of the hospitals (protocol numbers: ERC/2015/11/02 and LTH/ER/2016/01/254). Relevant biodata were recorded for each patient.

### Samples collection and bacterial identification

Aspirates were obtained from deep-seated abscesses while tissue samples were collected after washing the wound vigorously with sterile saline and debridement of the slough to exclude mere colonizers. Necrotic tissues were curetted into Anaerobic Basal Broth (Oxoid, Basingstoke, Hants, UK) for anaerobic culture. The samples were immediately transported to the laboratories and processed within 2 hours of sample collection by inoculating them onto a set of selective and non-selective media which were: 5% (v/v) sheep blood agar (BA: Oxoid, Basingstoke, Hants, UK), MacConkey agar (Oxoid, Basingstoke, Hants, UK), chocolate agar and Anaerobic Basal Agar (Oxoid, Basingstoke, Hants, UK) supplemented with 5% (v/v) laked sheep blood, Vit K1 (1μg/ml), L-cysteine hydrochloride (5 μg/ml) and gentamicin (100 μg/ml) (GBA).

A set of inoculated plain Blood agar and MacConkey agar were incubated in air at 37°C for 24 h, chocolate agar in CO_2_ at 37°C for 24 h. Another set of inoculated plain GBA, as well as GBA with kanamycin (75 μg/l) and vancomycin (5 μg/l) supplements, was incubated under atmospheric condition made up of 80% H_2_, 10% CO_2_, 10% N_2_ for 48 h and extended for 5 days if necessary: anaerobiosis was achieved using a Bactron Anaerobic Chamber (SHEL LAB, Cornelius, USA). Representative colonies were identified by colonial morphology, Gram staining characteristics and conventional biochemical tests using Microbact™ GNB 24E (Oxoid, Basingstoke, Hants, UK) and RapID™ STR (Remel, Lexena, USA) for facultative/aerobic Gram-negative bacilli and *Streptococcus* spp. respectively. *Staphylococcus* spp. was identified with catalase and coagulase tests. The obligate anaerobes were identified by RapID™ ANA II (Remel, Lexena, USA). Quality control strains, *Staphylococcus aureus* ATCC 25923, *Escherichia coli* ATCC 25922, *Bacteroides fragilis* ATCC 25285 and *Peptostreptococcus anaerobius* ATCC 27337, were used to assess the quality of the media and identification systems.

### Antibiotic susceptibility test

Antibiotic susceptibility testing for aerobic and facultative anaerobes was performed by modified Kirby-Bauer disk diffusion technique as recommended by Clinical and Laboratory Standards Institute (CLSI)^8^ using *Escherichia coli* ATCC 25922 and *S. aureus* ATCC 25923 as control strains. Discrete colonies were emulsified in sterile saline to match 0.5 McFaland turbidity standard from where confluence inocula were made on Mueller-Hinton agar (MHA) with sterile cotton swab. The swabbed MHA were allowed to dry at room temperature and a set of six antibiotics discs were placed evenly on each of them. After 18 - 24 hours of incubation, the diameter of the zone of inhibition around each antibiotic disc was measured, recorded and interpreted as “sensitive”, “intermediate” or “resistant” in accordance with CLSI guidelines.^8^ Isolates with intermediate sensitivity were regarded as “resistant”.

Extended-spectrum β-lactamase production was determined among Enterobacteriaceae and other GNB that have shown reduced susceptibility to at least one third generation cephalosporin or aztreonam by combination disc method according to CLSI guidelines.^8^ Gram-negative bacilli with intermediate sensitivity or resistance to one or more carbapenems were tested for production of carbapenemases by the Modified Hodge test (MHT) and interpreted by CLSI guidelines.^8^ Methicillin-resistant *S. aureus* (MRSA) was detected by disc diffusion test, using cefoxitin disc (30 μg) on Mueller-Hinton agar according to CLSI guidelines.^8^

Organisms that were phenotypically multidrug-resistant (MDR), including ESBL-producing GNB, carbapenem-resistant GNB and MRSA, were further tested for resistance determining genes using PCR-based protocols with specific oligonucleotide primers (Table 1) and template DNA of the bacteria extracted by boiling method.^9^ Electrophoresis of each PCR product (5μl) was carried out in 1.5% (w/v) Agarose gel (Biomatik, Ontario, Canada) in 1X Tris-Acetate-EDTA (TAE) buffer for 45 minutes. The size of amplified products was estimated using 100bp molecular weight marker (100 – 1200bp).

**Table 1:**
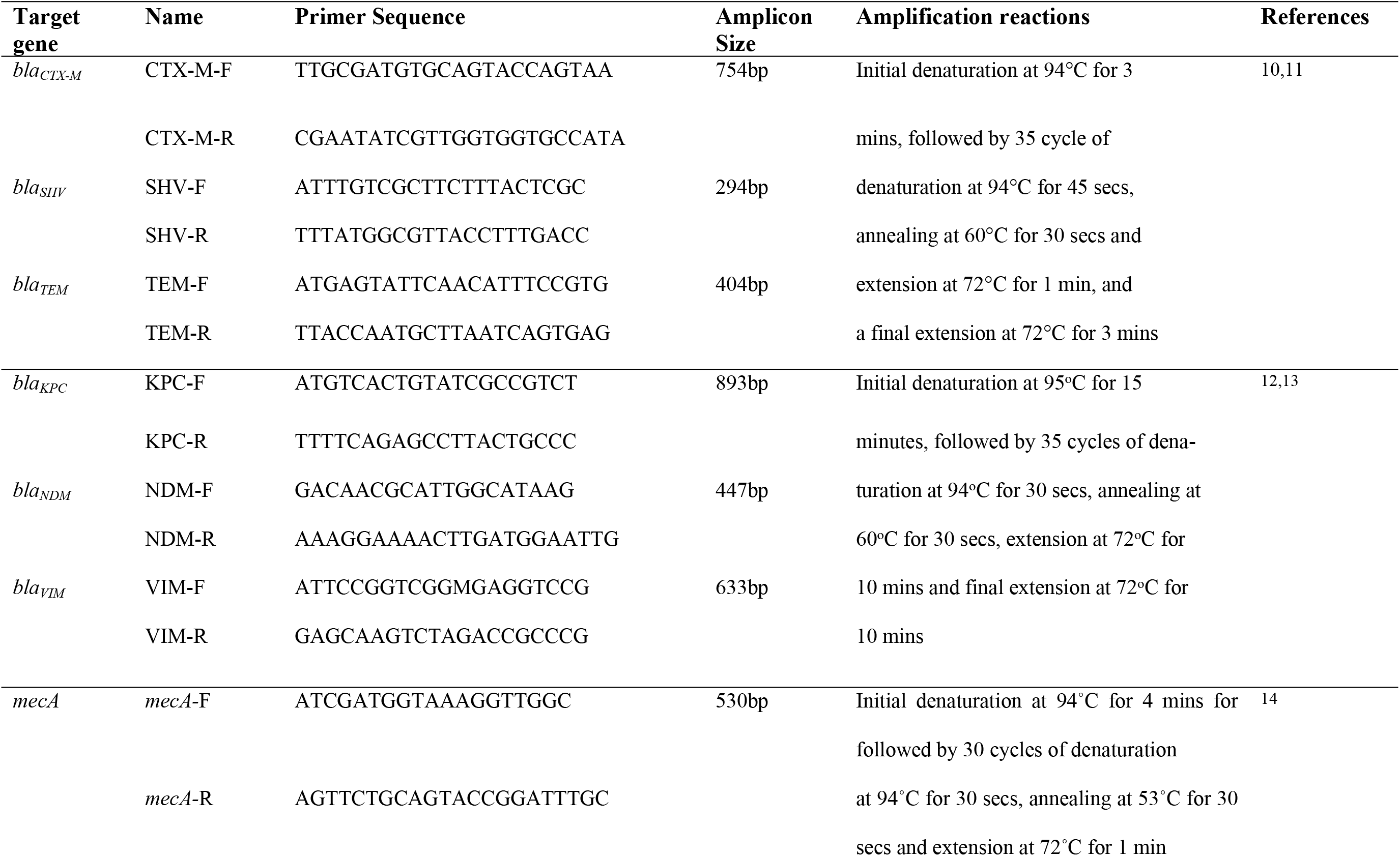
Oligonucleotides Primers and Amplification Reactions for Targeted Resistance Genes.

### Statistical analysis

Data analysis was performed with Statistical Package for Social Sciences (SPSS) version 20. Comparison of mean values was done using the Student’s t test for continuous and chi-square test for categorical variables. Risk factors for infection of diabetic foot by MDR organisms among were identified by logistic regression analysis. A p-value of 0.05 was considered to be statistically significant.

## Results

Ninety patients (53 males and 37 females) presented with 93 IDFUs during the 11-month study period. The patients ranged between 18 and 85 years (mean, 54.7 ± 12.8 years) of age. Of the 93 cases of foot infections, 70 (75.3%) were hospitalized patients and 56 (60%) had lasted for a duration of at least one month. Sixty-six (70.1%) of the ulcers were categorized as at least Wagner’s grade 3. The vast majority (n= 74; 82.2%) of the patients used antibiotics in the month before presentation at the facilities and over 90% (n= 84) had taken antibiotics before wound samples were obtained (Table 2).

**Table 2:**
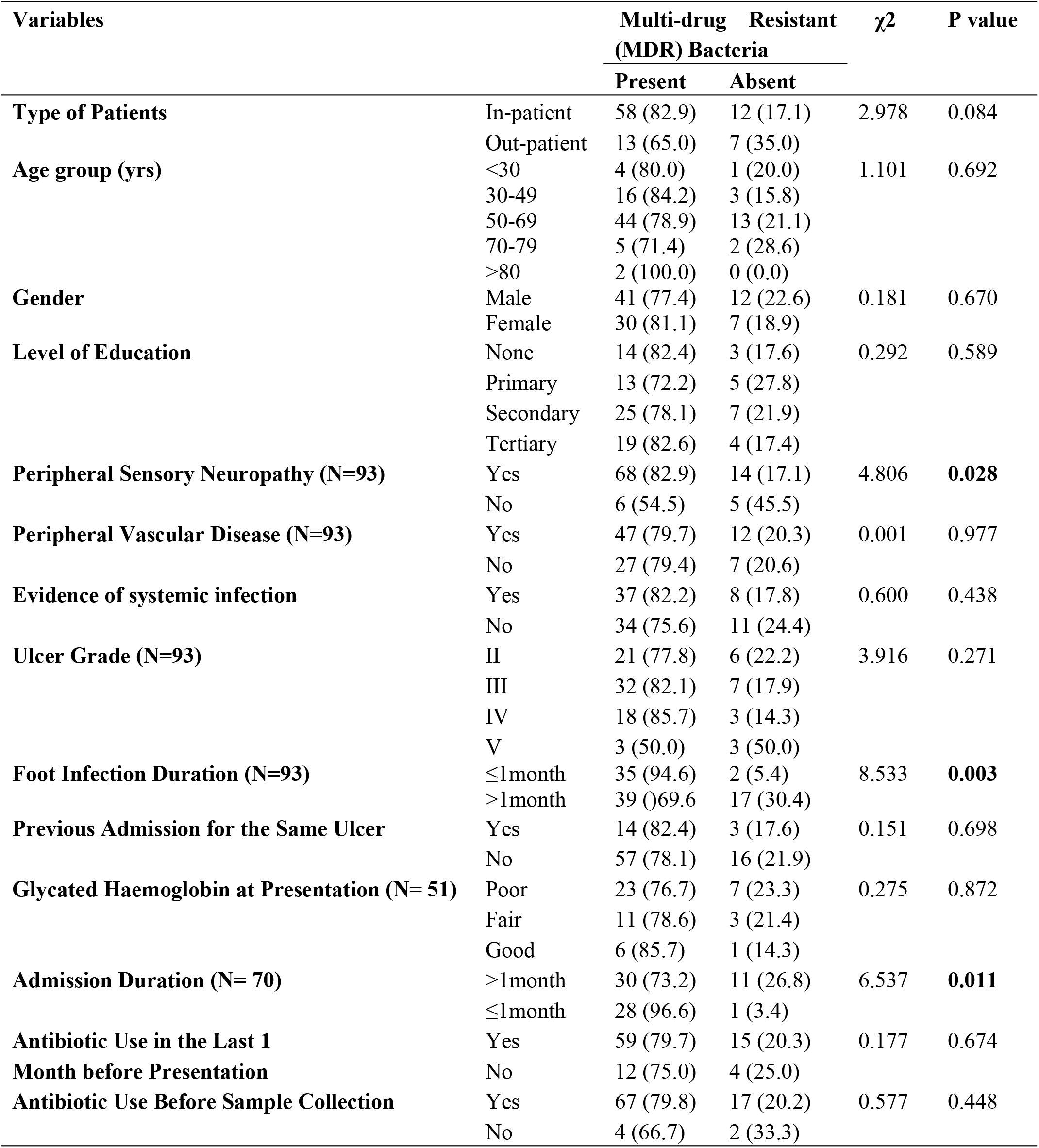
Association between Clinical/Socio-Demographic Variables of IDFU and MDR Bacteria (N= 90 unless otherwise stated)

| Variables | | Multi-drug Resistant (MDR) Bacteria | | $\chi^2$ | P value |
| --- | --- | --- | --- | --- | --- |
|  |  | Present | Absent |  |  |
| Type of Patients | In-patient | 58 (82.9) | 12 (17.1) | 2.978 | 0.084 |
|  | Out-patient | 13 (65.0) | 7 (35.0) |  |  |
| Age group (yrs) | <30 | 4 (80.0) | 1 (20.0) | 1.101 | 0.692 |
|  | 30-49 | 16 (84.2) | 3 (15.8) |  |  |
|  | 50-69 | 44 (78.9) | 13 (21.1) |  |  |
|  | 70-79 | 5 (71.4) | 2 (28.6) |  |  |
|  | >80 | 2 (100.0) | 0 (0.0) |  |  |
| Gender | Male | 41 (77.4) | 12 (22.6) | 0.181 | 0.670 |
|  | Female | 30 (81.1) | 7 (18.9) |  |  |
| Level of Education | None | 14 (82.4) | 3 (17.6) | 0.292 | 0.589 |
|  | Primary | 13 (72.2) | 5 (27.8) |  |  |
|  | Secondary | 25 (78.1) | 7 (21.9) |  |  |
|  | Tertiary | 19 (82.6) | 4 (17.4) |  |  |
| Peripheral Sensory Neuropathy (N=93) | Yes | 68 (82.9) | 14 (17.1) | 4.806 | **0.028** |
|  | No | 6 (54.5) | 5 (45.5) |  |  |
| Peripheral Vascular Disease (N=93) | Yes | 47 (79.7) | 12 (20.3) | 0.001 | 0.977 |
|  | No | 27 (79.4) | 7 (20.6) |  |  |
| Evidence of systemic infection | Yes | 37 (82.2) | 8 (17.8) | 0.600 | 0.438 |
|  | No | 34 (75.6) | 11 (24.4) |  |  |
| Ulcer Grade (N=93) | II | 21 (77.8) | 6 (22.2) | 3.916 | 0.271 |
|  | III | 32 (82.1) | 7 (17.9) |  |  |
|  | IV | 18 (85.7) | 3 (14.3) |  |  |
|  | V | 3 (50.0) | 3 (50.0) |  |  |
| Foot Infection Duration (N=93) | ≤1month | 35 (94.6) | 2 (5.4) | 8.533 | **0.003** |
|  | >1month | 39 (69.6) | 17 (30.4) |  |  |
| Previous Admission for the Same Ulcer | Yes | 14 (82.4) | 3 (17.6) | 0.151 | 0.698 |
|  | No | 57 (78.1) | 16 (21.9) |  |  |
| Glycated Haemoglobin at Presentation (N= 51) | Poor | 23 (76.7) | 7 (23.3) | 0.275 | 0.872 |
|  | Fair | 11 (78.6) | 3 (21.4) |  |  |
|  | Good | 6 (85.7) | 1 (14.3) |  |  |
| Admission Duration (N= 70) | >1month | 30 (73.2) | 11 (26.8) | 6.537 | **0.011** |
|  | ≤1month | 28 (96.6) | 1 (3.4) |  |  |
| Antibiotic Use in the Last 1 Month before Presentation | Yes | 59 (79.7) | 15 (20.3) | 0.177 | 0.674 |
|  | No | 12 (75.0) | 4 (25.0) |  |  |
| Antibiotic Use Before Sample Collection | Yes | 67 (79.8) | 17 (20.2) | 0.577 | 0.448 |
|  | No | 4 (66.7) | 2 (33.3) |  |  |

Results further showed a total of 218 organisms were isolated from the 93 specimens examined with an average of 2.34 organisms per sample. Of the organisms, 129 (59.2%) were Gram-negative aerobic bacilli, 59 (27.1%) Gram-positive aerobic cocci and 29 (13.2%) anaerobic bacteria; only one (0.5%) was yeast. *S. aureus*, 41 (18.8%) was the single most common organism followed by *E. coli,* 23 (10.6%) and *Pseudomonas aeruginosa,* 20 (9.2%). Others included *Klebsiella* spp. (19; 8.7%), *Citrobacter* spp. (19; 8.7%), *Enterococcus* spp. (13; 6%), *Enterobacter* spp. (11; 5.1%), *Proteus mirabilis* (10; 4.6%) and *Acinetobacter* spp. (9; 4.1%). On the other hand, the predominant anaerobic bacteria were *Bacteroides* spp. (7; 3.2%) and *Peptostreptococcus anaerobius* (6; 2.3%) as shown in Table 3.

**Table 3:**
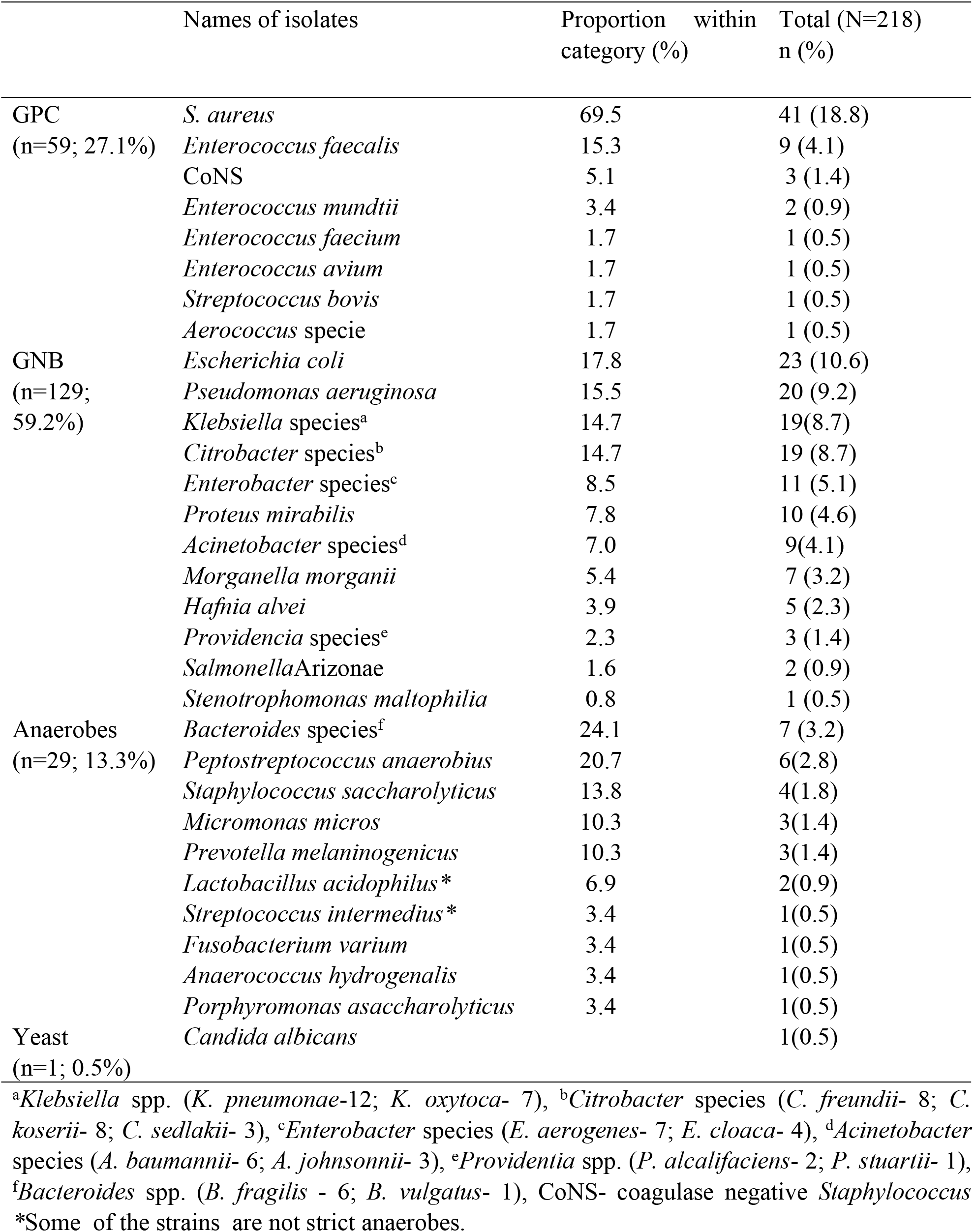
Bacterial etiological agents of infected diabetic foot ulcers.

Gram-positive bacteria were highly resistant to co-trimoxazole (69.5%), penicillin G (66.1%) and gentamicin (40.1%) but minimally resistant to piperacillin/tazobactam (6.8%) and amikacin (10.2%). On the other hand, Gram-negative bacteria were highly resistant to the tested third generation cephalosporins which included ceftriaxone (56%), cefotaxime (55%) and ceftazidime (48.1%); cotrimoxazole (89%), gentamicin (54.3%) and ciprofloxacin (54.3%). Low rates of resistance were found to ertapenem (6.4%), piperacillin/tazobactam (9.3%) and amikacin (12.4%).

Analysis of antibacterial resistance profiles of the organisms showed that of the188 aerobic isolates, 121 (64.4%) were multidrug-resistant (MDR), being resistant to one or more agents in at least three antibiotic classes (Table 4). Further analysis of specific MDR phenotypes showed that 13 (31.7%) of *S. aureus* were methicillin-resistant (MRSA), while 43 (33.3%) and 10 (7.8%) of Gram-negative bacteria were ESBL-producing and carbapenem-resistant respectively (Table 5). Ten (76.9%) of the methicillin-resistant *S. aureus* isolates harbored *mec*A gene and 37 (86.1%) of the 43 ESBL-producing phenotypes harbored at least one of the ESBL-determining genes sought with the most common being *bla*_CTX-M_ (30; 81.1%). Others were *bla*_TEM_ (27; 72.8%) and *bla*_SHV_ (8; 21.6%). Multiple genes were detected in 72.1% of ESBL-producing GNB (n= 31). Carbapenemase determinants were detected in four (40%) of the 10 carbapenem-resistant organisms; they were *bla*_VIM_ (3; 30%), *bla*_KPC_ (2; 20%) and *bla*_NDM_ (1; 10%).

**Table 4:** Prevalence of Multidrug-resistant bacterial isolates in IDFU.

| **BACTERIA ISOLATES (N)** | **MDR (n)** | **PREVALENCE (%)** |
| --- | --- | --- |
| **Gram positive cocci (59)** | **33** | **59.9** |
| *Staphylococcus aureus* (41) | 20 | 48.8 |
| *Enterococcus* spp. (13) | 11 | 84.6 |
| CoNS (3) | 0 | 0.0 |
| *Streptococcus bovis* (1) | 1 | 100.0 |
| *Aerococcus* spp. (1) | 1 | 100.0 |
| **Gram negative bacilli (129)** | **88** | **68.2** |
| *Escherichia coli* (23) | 16 | 69.6 |
| *Pseudomonas aeruginosa* (20) | 10 | 50.0 |
| *Klebsiella* species (19) | 13 | 68.4 |
| *Citrobacter* species (19) | 14 | 73.7 |
| *Enterobacter* species (11) | 9 | 81.8 |
| *Proteus mirabilis* (10) | 6 | 60.0 |
| *Acinetobacter* species (9) | 8 | 88.9 |
| *Morganella morganii* (7) | 4 | 57.1 |
| *Hafnia alvei* (5) | 4 | 80.0 |
| *Providencia* species (3) | 2 | 66.7 |
| *Salmonella* Arizonae (2) | 1 | 50.0 |
| *Stenotrophomonas maltophilia* (1) | 1 | 100.0 |
| **TOTAL (188)** | **121** | **64.4** |

MDR – multidrug-resistant, CoNS- coagulase negative *Staphylococcus*

**Table 5:** Types of multidrug-resistant bacteria in IDFU cases.

| **BACTERIA ISOLATES (N=188)** | **MRSA/MRCoNS n (%)** | **VRSA/VRE n (%)** | **ESBL n (%)** | **AmpC n (%)** | **Carbapenem Resistance n (%)** |
| --- | --- | --- | --- | --- | --- |
| **GPC (N=59)** |  |  |  |  |  |
| *S. aureus* (41) | 13 (31.7) | 0 (0.0) |  |  |  |
| *Enterococcus* spp (13) |  | 0 (0.0) |  |  |  |
| *Streptococcus bovis* (1) |  |  |  |  |  |
| *Aerococcus* sp (1) |  |  |  |  |  |
| CoNS (3) | 0 (0.0) |  |  |  |  |
| **GNB (N=129)** |  |  | **43 (33.3)** | **4 (3.1)** | **10 (7.8)** |
| *Escherichia coli* (23) |  |  | 10 (43.5) | 2 (8.7) |  |
| *Klebsiella* spp (19) |  |  | 10 (52.6) |  |  |
| *Citrobacter* spp (19) |  |  | 10 (52.6) | 1 (5.3) |  |
| *Enterobacter* spp (11) |  |  | 6 (54.6) |  |  |
| *P. aeruginosa* (20) |  |  |  |  | 3 (15.0) |
| *Acinetobacter* spp (9) |  |  |  |  | 4 (44.4)* |
| *M. morganii* (7) |  |  | 2 (28.8) | 1 (14.3) | 1 (14.3) |
| *P. mirabilis* (10) |  |  | 2 (20.0) |  |  |
| *Hafnia alvei* (5) |  |  | 1 (20.0) |  | 2 (40.0)*** |
| *S. maltophilia* (1) |  |  | 1 (100.0) |  |  |
| *Providencia* spp (3) |  |  | 1 (33.3)** |  |  |

\*Specifically *A. baumannii*, \*\*Specifically *P. alcalifaciens*, \*\*\*Carbapenemase-producing (Modified Hodge Test Positive),
MRSA – methicillin-resistant *Staphylococcus aureus*, MRCoNS – methicillin-resistant coagulase negative *Staphylococcus*, VRSA – vancomycin-resistant *Staphylococcus aureus*, VRE – vancomycin-resistant *Enterococcus*, ESBL – extended-spectrum β-lactamase, AmpC – AmpC β-lactamase. GPC – Gram-positive cocci, GNB – Gram-negative bacilli

Significant factors associated with presence of MDR organisms in diabetic foot infections included peripheral sensory neuropathy, foot infection duration >1 month and admission duration >1 month (Table 2). Further analysis with logistic regression however identified only peripheral neuropathy (r= 4.05, p= 0.042) and foot infection duration >1 month (r= 7.63, p= 0.015) as the predisposing factors for acquisition of multidrug-resistant bacteria among patients with diabetic foot infection.

## Discussion

Infected diabetic ulcers continue to be a polymicrobial infection involving aerobic as well as obligate anaerobic organisms. All IDFUs studied in this series have at least an average of two different bacteria implicated in the disease. Gram-negative bacteria including *E. coli*, *Pseudomonas aeruginosa*, *Klebsiella* species and *Enterobacter* species predominate, which reflects the long-standing nature of the infections, itself a consequence of poor health-seeking behavior in this part of this world.^2^ Furthermore, a wide range of anaerobic bacteria primarily *Bacteroides* species and *Peptostreptococcus anaerobius* are important agents of the infections and were isolated from a third of the cases. This suggests infections that are chronic and below the superficial layers of the skin.^15^

Antibiotic resistance remains a huge problem among diabetic foot ulcer infections; it worsens prognosis and makes treatment outcomes poor.^16^ Seventy four (80%) of the 93 IDFU cases in this series harbour one or more MDR bacteria, largely attributable to inappropriate antibiotics use and unrestricted access to antimicrobial drugs in many low- and middle-income countries.^17^ This contrasts with several studies in high-income countries including France with low prevalence of MDR bacteria among patients with IDFU.^16,18^ A wide spectrum of aerobic and facultative anaerobic bacteria are found to be multidrug-resistant in this study, comparable to findings elsewhere in Africa and Asia.^19,20^

A third of the *S. aureus* isolates were methicillin-resistant (MRSA). Though prevalence of MRSA appears to be rising in Africa, most of the countries have rates lower than 50%.^21^ This study also revealed that *mecA* was detected in 77% of the MRSA and this is similar to the observation of Chaudhry *et al.* who detected the gene in 20 (84%) of the 25 phenotypically confirmed MRSA isolates.^22^ That there are MRSA that lack *mec*A gene may be on account of *mecC,* a variant of *mecA* discovered in 2011 as well as other mutations of penicillin-binding proteins as alternate mechanisms of penicillin resistance.^23^

Extended-spectrum β-lactamases were produced by 33.3% of all Gram-negative bacilli isolated and all the organisms except two were members of the family *Enterobacteriaceae*. The leading bacterial hosts producing ESBLs were *E. coli, Klebsiella* and *Citrobacte*r species. Published rates of ESBL-producing bacteria vary widely across countries in Africa; as high as 96% in Mali to 0.3% in South Africa.^24^ have all been reported. The most prevalent ESBL type was the CTX-M which has been reported as the most predominant variant worldwide.^25^ In this study, only 10 (7.8%) of the Gram-negative bacteria were resistant to the carbapenems. Carbapenem resistance determining genes were present in *Acinetobacter baumannii*, *Hafnia alvei* and *Morganella morganii*. Carbapenems are drugs of last resort in the treatment of resistant Gram-negative bacilli infections and with variable and increasing rates of resistance being reported.^26,27^

Independent risk factors for acquisition of MDR bacteria found in our study are peripheral sensory neuropathy and foot infection duration > 1 month. This is similar to reports among IDFU cases from India.^28,29^ Other authors also documented prolonged duration of wound infection as a predictor of infection of diabetic foot ulcers with MDR bacteria.^30,31^ Contrary findings have however been documented from other studies in China, Iran and Portugal.^18,20,32^ Our findings is also discordant with the report of Noor *et al.* which established that ulcer size is a risk factor for infection by multidrug-resistant organisms.^31^ This study also observed a significant association between presence of multidrug-resistant bacteria in IDFU and long duration of hospitalization (>1 months) similar to previously documented reports by another author in Turkey.^33^ We did not find any socio-demographic factors that were significantly associated with occurrence of MDR IDFU in our study as with other reports.^28–30^ However, on the contrary to our finding, Trivedi *et al.* in the United States noted smoking as an independent risk factor for multidrug-resistant foot wound infection.^34^

## Conclusion

The spectrum of agents causing IDFU is wide and includes numerous species of aerobic and anaerobic bacteria. There is a high prevalence of MDR aerobic bacteria among them which poses a great limitation to effective treatment of cases. Improved health seeking attitude and timely antibiotic coverage for MDR bacteria will therefore mitigate IDFU in our environment.

## Abbreviations

IDFU: Infected diabetic foot ulcer
SPSS: Statistical Package for Social Sciences
GNB: Gram-negative bacilli
GPC: Gram-positive cocci
MDR: multidrug resistant
CLSI: Clinical and Laboratory Standards Institute
MHA: Mueller Hinton agar
MRSA: methicillin-resistant *Staphylococcus aureus*
ESBL: extended-spectrum β-lactamase

## Declarations

### Ethics approval and consent to participate

Ethical approvals for this study were granted by the Ethics and Research Committees of the Obafemi Awolowo University Teaching Hospitals Complex and Ladoke Akintola University College of Technology with protocol numbers ERC/2015/11/02 and LTH/ER/2016/01/254 respectively.

### Consent for publication

Not applicable.

### Availability of data and material

The datasets used and analysed during the current study are available from the corresponding author on reasonable request.

### Authors’ contributions

ATA and AOA conceived and designed the study. BAK and VOR contributed to the design of the study. ATA and AOA conducted laboratory experiments. ATA and AOA analyzed the data. ATA, AOA and VOR wrote the final report. All authors reviewed and approved the final report.

## Acknowledgements

The authors are grateful to Faith Ayobami for assistance in data management and Babatunde Odetoyin for technical assistance in the Molecular Biology Laboratory.

## References

1. Uckay I, Gariani K, Pakaty Z, Lipsky BA. Diabetic foot infections: state of the art. Diabetes, Obes Metab 2014; 6(4): 305–16.

2. Lipsky BA, Berendt AR, Cornia PB, Pile JC, Peters EJG, Armstrong DG, et al. 2012 Infectious Diseases Society of America clinical practice guideline for the diagnosis and treatment of diabetic foot infections. Clin Infect Dis 2012; 54(12): 132–73

3. Zhang P, Lu J, Jing Y, Tang S, Zhu D, Bi Y. Global epidemiology of diabetic foot ulceration: a systematic review and meta-analysis. Ann Med. 2017r; 49(2):106–16. doi: 10.1080/07853890.2016.1231932.

4. Atun R and Gale EA. The challenge of diabetes in sub-Saharan Africa. Lancet Diabetes Endocrinol. 2015 Sep; 3(9): 675–7. doi: 10.1016/S2213-8587(15)00236-3.

5. Jia L, Parker CN, Parker TJ, et al. Incidence and risk factors for developing infection in patients presenting with uninfected diabetic foot ulcers. Jan Y-K, ed. PLoS ONE. 2017; 12(5): e0177916. doi: 10.1371/journal.pone.0177916.

6. Ventola CL. The Antibiotic Resistance Crisis: Part 1: Causes and Threats. Pharmacy and Therapeutics 2015; 40(4) :277–83.

7. Laxminarayan R, Matsoso P, Pant S, Brower C, Røttingen JA, Klugman K, Davies S. Access to effective antimicrobials: a worldwide challenge. Lancet 2016; 387(10014): 168–75. doi: 10.1016/S0140-6736(15)00474-2.

8. CLSI. Performance Standards for Antimicrobial Susceptibility Testing; Twenty-Fifth Informational Supplement. CLSI document M100-S25. Wayne, PA: Clinical and Laboratory Standards Institute; 2015.

9. Dashti AA, Jadaon MM, Abdulsamad AM, Dashti HM. Heat Treatment of Bacteria: A Simple Method of DNA Extraction for Molecular Techniques. Kuwait Med J. 2009; 41(2): 117–22.

10. Sidjabat HE, Paterson DL, Adams-Haduch JM, Ewan L, Pasculle AW, Muto CA, Tian GB, Doi Y. Molecular Epidemiology of CTX-M-producing Escherichia coli isolates at a Tertiary Medical Center in Western Pennsylvania. Antimicrobial Agents Chemother 2009; 53(11): 4733–9.

11. Monstein HJ, Ostholm-Balkhed A, Nilsson MV, Dornbusch NL. Multiplex PCR amplification assay for rapid detection of SHV, TEM and CTX-M genes in Enterobateriaceae. APMIS 2007; 115: 1400–8.

12. Pasanen T, Koskela S, Mero S, Tarkka E, Tissari P, Vaara M, et al. Rapid Molecular Characterization of *Acinetobacter baumannii* Clones with rep-PCR and Evaluation of Carbapenemase Genes by New Multiplex PCR in Hospital District of Helsinki and Uusimaa. PLoS ONE 2014; 9(1): e85854.

13. Zhao S, Jang D, Xu P, Xang Y, Shi H, Cao H et al. An Investigation of Drug-Resistant Acinetobacter baumannii Infection in a Comprehensive Hospital in East India. Ann Clin Microbiol and Antimicrob. 2015; 14:7. doi: 10.1186/s12941-015-0066-4.

14. Sajith Khan AK, J Shetty, Preetha Lakshmi Sarayu Y, Anandi Chidambaram and Ramesh Ranganathan. Detection of *mecA* genes of Methicillin-Resistant *Staphylococcus aureus* by Polymerase Chain Reaction. Inter J. Health Rehab Sci. 2012; 1: 64–8.

15. Mendes JJ, Nerves J. Diabetic foot infections: current diagnosis and treatment. The J. Diab Foot Compl 2012; 4(1): 26–45.

16. Richard JL, Sotto A, Jourdan N, Combescure C, Vannereau D, Rodier M, et al. Risk factors and healing impact of multidrug-resistant bacteria in diabetic foot ulcers. Diab Metab 2008; 34(2): 363–9.

17. Omolase CO, Adeleke OE, Afolabi AO, Afolabi OT. Self medication amongst general out-patients in a Nigerian community hospital. Annals of Ibadan Postgrad Med 2007: 5(2): 64–7.

18. Mendes JJ, Marques-Costa A, Vilela C, Neves J, Candeias N, Cavaco-Silva P, et al. Clinical and bacteriological survey of diabetic foot infections in Lisbon. Diab Res and Clin Prac 2012; 95(1): 153–61.

19. Dwedar R, Ismail DK, Abdulbaky A. Lecturer Diabetic foot Infection: Microbiological Causes with Special Reference to their Antibiotic Resistance Pattern. Egyptian J. Med Microbiol 2015; 24(3): 95–102.

20. Amini M, Davati A, Piri M. Determination of the Resistance Pattern of Prevalent Aerobic Bacterial Infections of Diabetic Foot Ulcer. Iranian J. Pathol 2013; 8(1): 21–6.

21. Falagas ME, Karageorgopoulos DE, Leptidis J, Korbila IP. MRSA in Africa: Filling the global map of antimicrobial resistance. PLoS One 2013; 8(7): e68024. doi: 10.1371/journal.pone.0068024.

22. Chaudhry WN, Badar R, Jamal M, Jeong J, Zafar J, Andleeb S. Clinico‑microbiological study and antibiotic resistance profile of *mecA* and ESBL gene prevalence in patients with diabetic foot infections. Experimental and Therapeutic Medicine 2016; 11: 1031–38.

23. Shore AC, Deasy EC, Slickers P, Brennan G, O’Connell B, Monecke S,Ehricht R, Coleman DC. Detection of staphylococcal cassette chromosome mec type XI carrying highly divergent mecA, mecI, mecR1, blaZ,and ccr genes in human clinical isolates of clonal complex 130 methicillin-resistant *Staphylococcus aureus*. Antimicrob Agents Chemother 2011: 55: 3765–73. https://doi.org/10.1128/AAC.00187-11.

24. Motta RN, Oliveira MM, Magalhães PSF, Dias AM, Aragão LP, Forti A.C et al. Plasmid-mediated extended-spectrum β-lactamase-producing strains of Enterobacteriaceae isolated from diabetes foot infections in a Brazilian diabetic center. Braz J. Infect Dis 2003; 7(2): 129–34.

25. Shaikh S, Fatima J, Shakil S, Rizvi SMD, Kamal MA. Antibiotic resistance and extended spectrum beta-lactamases: Types, epidemiology and treatment. Saudi J. Biol Sci. 2015; 22 (1): 90–101. doi: 10.1016/j.sjbs.2014.08.002.

26. Mohammed Y, Zailani SB, Onipede AO. Characterization of KPC, NDM and VIM Type Carbapenem Resistance Enterobacteriaceae from North Eastern, Nigeria. J. Biosciences and Medicines 2015; 3: 100–107.

27. Mushi MF, Mshana SE, Imirzalioglu C, Bwanga F. Carbapenemase Genes among Multidrug Resistant Gram Negative Clinical Isolates from a Tertiary Hospital in Mwanza, Tanzania. BioMed Res Inter 2014; 2014: 303104. http://dx.doi.org/10.1155/2014/303104.

28. Gadepalli R, Dhawan B, Sreenivas V, Kapil A, Amini AC, Chaudhry RA. Clinico-microbiological study of diabetic foot ulcers in an Indian Tertiary Care Hospital. Diabetes Care 2006; 29(8): 1727–32.

29. Zubair M, Abida M, Jamal A. Clinicobacteriology and risk factors for the diabetic foot infection with multidrug resistant microorganisms in North India. Biol Med 2010; 2(4): 22–34.

30. Hartemann-Heurtier A, Robert J, Jacqueminet S, Van GH, Golmard JL, Jarlier, et al. Diabetic foot ulcer and multidrug-resistant organisms: risk factors and impact. Diabetic Medicine 2004; 21: 710–15.

31. Noor S, Borse AG, Ozair M, Raghav A, Parwez I, Ahmad J. Inflammatory markers as risk factors for infection with multidrug-resistant microbes in diabetic foot subjects. Foot (Edinb). 2017; 32: 44–8. doi: 10.1016/j.foot.2017.05.001

32. Ji X, Jin P, Chu Y, Feng S, Wang P. Clinical characteristics and risk factors of diabetic foot ulcer with multidrug-resistant organism infection. Inter J. of Lower Extrem Wounds 2014; 13(1): 64–71.

33. Kandemir Ö, Akbay E, Şahin E, Milcan A, Gen R. Risk factors for infection of the diabetic foot with multi-antibiotic resistant microorganisms. J. of Infect 2007; 54(5): 439–45.

34. Trivedi U, Parameswaran S, Armstrong A, Burgueno-Vega D, Griswold J, Dissanaike S, Rumbaugh KP. Prevalence of Multiple Antibiotic Resistant Infections in Diabetic versus Nondiabetic Wounds. J Pathog. 2014;2014:173053. doi: 10.1155/2014/173053.

